# Myo1f has an essential role in γδT intraepithelial lymphocyte adhesion and migration

**DOI:** 10.1101/2022.08.30.505932

**Authors:** Irving Ulises Martínez-Vargas, Maria Elena Sánchez-Bello, Carlos Emilio Miguel-Rodríguez, Felipe Hernández-Cázares, Leopoldo Santos-Argumedo, Patricia Talamás Rohana

## Abstract

γδT intraepithelial lymphocyte represents up to 60% of the small intestine intraepithelial compartment. They are highly migrating cells and constantly interact with the epithelial cell layer and lamina propria cells. This migratory phenotype is related to the homeostasis of the small intestine, the control of bacterial and parasitic infections, and the epithelial shedding induced by LPS. Here, we demonstrate that Myo1f participates in the adhesion and migration of intraepithelial lymphocytes. Using long-tailed class I myosins KO mice, we identified the requirement of Myo1f for their migration to the small intestine intraepithelial compartment. The absence of Myo1f affects intraepithelial lymphocytes’ homing due to reduced CCR9 and a4b7 expression. *In vitro*, we confirm that adhesion to integrin ligands and CCL25-dependent and independent migration of intraepithelial lymphocytes are Myo1f-dependent. Mechanistically, Myo1f deficiency prevents correct chemokine receptor and integrin polarization, leading to reduced tyrosine phosphorylation and having consequences in cell signaling. Overall, we demonstrate that Myo1f has an essential role in the adhesion and migration in γδT intraepithelial lymphocytes.

**SUMMARY STATEMENT:** The adhesion and migration of γδT intraepithelial lymphocytes are regulated by the motor protein Myosin 1f through controlling the expression of integrins and the chemokine receptor CCR9.

## INTRODUCTION

Intraepithelial lymphocytes are cells that reside in the epithelial cell layer of the mucosa. Intestinal intraepithelial lymphocytes are mainly T lymphocytes, either αβT or γδT. These cells are sub-classified into conventional T lymphocytes, CD4 and CD8αβ αβT, and unconventional CD8αα αβT and γδT (Cheroutre et al., 2011). The most unconventional T cells are γδT lymphocytes which constitute up to 60% of the intraepithelial lymphocytes in the small intestine (Goodman and Lefrancois, 1988; Boll et al., 1995; Bonneville et al., 1988). These cells were described as anchored cells in the intestinal epithelium (Darlington and Rogers, 1996). However, seminal works have shown that intestinal intraepithelial γδT lymphocytes migrate between the intestinal epithelium and the basal lamina in the small intestine (Edelblum et al., 2012; Edelblum et al., 2015; van Konijnenburg et al., 2017; Sumida et al., 2017; Fischer et al., 2020). This migration was related to the homeostasis of the intestinal epithelium (Sumida et al., 2017), the control of infections by *Salmonella* and *Toxoplasma gondii* (Edelblum et al., 2015; van Konijnenburg et al., 2017), and the pathological shedding (Hu et al., 2022).

Intraepithelial lymphocyte homing depends mainly on CCR9 and α4β7 integrin (Wurbel et al., 2001; Uehara et al., 2002; Wagner et al., 1996). In addition, the interstitial migration of these cells depends on other molecules such as occludin (Edelblum et al., 2012), integrin αE (Schön et al., 1999), orphan receptors such as GRP18 (Wang et al., 2014) and GPR55 (Sumida et al., 2017), and IL-15 (Hu et al., 2018).

Class I myosins are monomeric myosins that link the actin cytoskeleton to the plasma membrane through their motor and pleckstrin homology (PH) domain, respectively (Krendel and Mooseker, 2005). These myosins are expressed in lymphoid and myeloid cells and have been related to cell adhesion, migration, and vesicular trafficking (Maravillas-Montero and Santos-Argumedo, 2012; Girón-Pérez et al., 2019). Among the class I myosins, Myo1e and Myo1f are two long-tailed class I myosins with two additional domains in their tail region. Both contain a proline-rich sequence (TH2) and an SH3 domain that recognizes proline-rich sequences, which allow protein-protein interactions (Krendel and Mooseker, 2005; Navinés-Ferrer and Martín, 2020).

Myo1e participates in neutrophils and B cells integrin mediated-adhesion and migration (Vadillo et al., 2019; Girón-Pérez et al., 2020) as well as spreading (Wenzel et al., 2015). Similarly, Myo1f is relevant for neutrophil migration (Kim et al., 2006; Salvermoser et al., 2018; Wang et al., 2019). Particularly, Myo1f has shown to be relevant for the β2 integrin expression in neutrophils (Kim et al., 2006), β1 and β7 integrins in mast cells (Navinés-Ferrer et al., 2019), and αVβ3 integrin in macrophages (Piedra-Quintero et al., 2019); presumably due to alterations in vesicular traffic (Kim et al., 2006). In this regard, Myo1f deficiency results in defects in IgE- and MRGPRX2-dependent mast cell degranulation and TNF secretion (Navinés-Ferrer et al., 2021). Mechanically, Myo1f is required for Cdc42 activation, and its deficiency leads to decreased cortical actin polymerization (Navinés-Ferrer et al., 2021). Additionally, Myo1f has emerged as an α-tubulin interacting protein relevant in the dynein-mediated Syk and CARD9 protein transport from the membrane to the cytoplasm during antifungal activation in macrophages (Sun et al., 2021). Finally, Myo1f regulates filopodia dynamics, increasing adhesion and migration (Hensel A. et al., 2022). Together, indicating the relevance of the role of Myo1f during the dynamics of the plasma membrane, cell adhesion and migration, and vesicular trafficking.

Here, we show that intraepithelial lymphocytes express Myo1e and Myo1f. Furthermore, the absence of Myo1e and Myo1f impacts the homing of intraepithelial γδT lymphocytes, but Myo1f shows a more significant impact. This deficiency was due to a reduction in the expression of intestinal homing receptors CCR9 and α4β7 integrin, having consequences on the 2D migration of γδT lymphocytes *in vitro*. Mechanistically, Myo1f was relevant for actin-mediated membrane protrusion formation during adhesion and migration. Its absence impacts CCR9 and integrin polarization and leads to tyrosine phosphorylation defects, which have signaling consequences.

## RESULTS

### Intraepithelial lymphocytes express long-tailed class I myosins

There is no evidence of class I myosin protein expression by intraepithelial lymphocytes. As a first approach, mRNA class I myosins expression was analyzed in intraepithelial, thymus, and spleen γδT lymphocytes with data from The Immunological Genome Project (https://www.immgen.org/). Heat map reveals that among class I myosins, Myo1e and Myo1f are expressed mainly by intraepithelial γδT lymphocytes, both Vγ7+ and Vγ7-. Myo1f was also expressed in activated spleen Vγ4+ and Vγ4-γδT lymphocytes but did not in resting spleen γδT lymphocytes. Mature thymic Vγ5 γδT lymphocytes also express Myo1f. Whereas the rest of class I myosins remain less represented in γδT lymphocytes (Fig. 1A). These bioinformatic data suggest that Myo1f is the main class I myosin expressed in γδT lymphocytes. Next, the expression at the protein level of Myo1e and Myo1f was confirmed by western blot. Myo1e and Myo1f are expressed in total IELs (Fig. 1B). Densitometric analysis confirmed that Myo1f is more abundant than Myo1e, as observed in western blot band thickness (Fig. 1C). Flow cytometry analysis of Myo1e and Myo1f expression confirmed the results obtained by western blot. Analysis of total IELs showed similar results in γδT and αβT IEL (Figs. 1D, S1A, B), whereas thymic and spleen γδT lymphocytes do not express Myo1e but do express Myo1f; however, in less proportion than IEL (Figs. S1C, D). Mean Fluorescence Intensity (MFI) analysis confirmed that Myo1f was expressed more than Myo1e in total IELs and γδT IEL (Fig. 1E). Thus, intraepithelial lymphocytes express long-tailed class I myosins.

**Figure 1.**
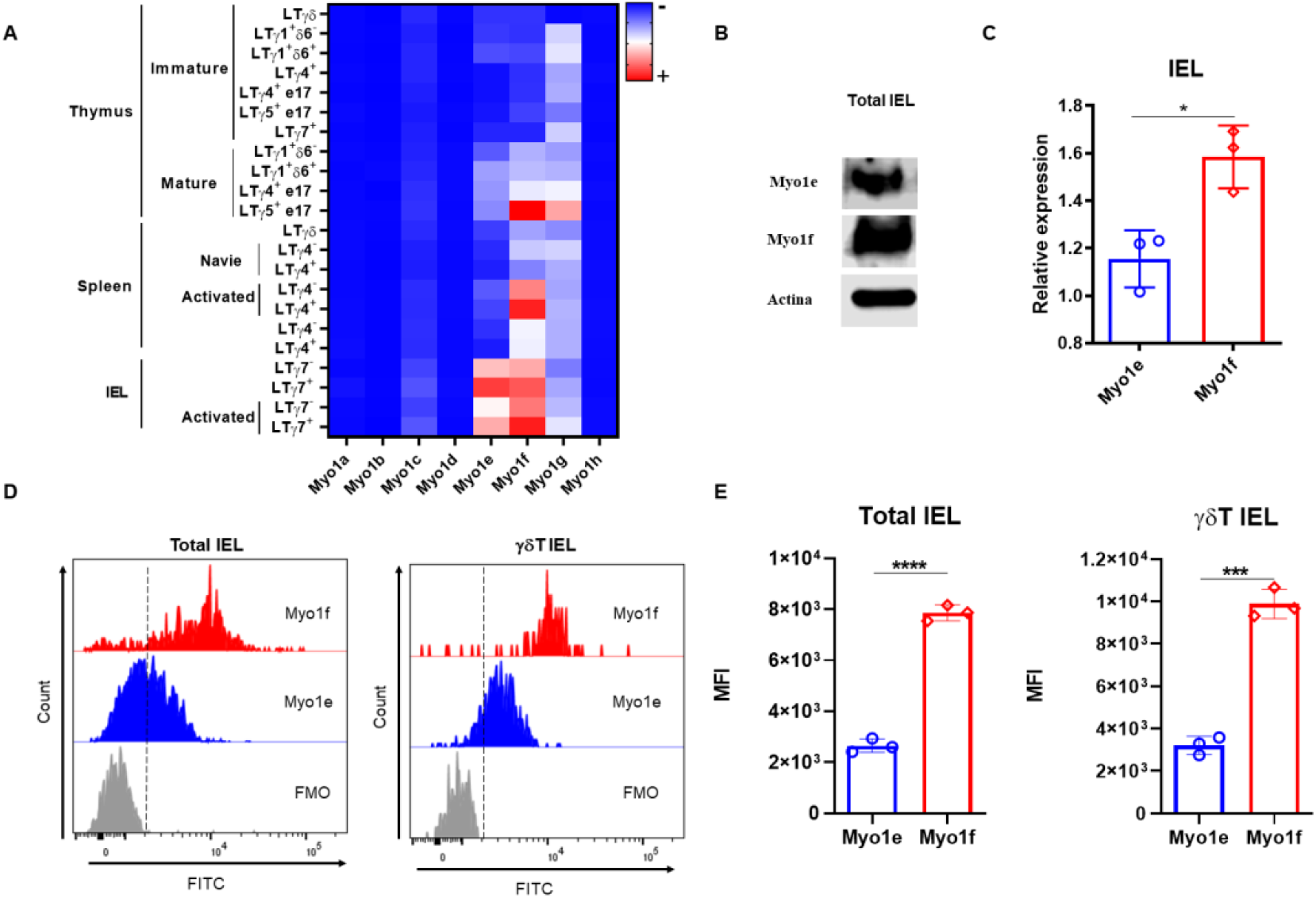
Expression of class I myosins in IEL. **A)** Heat map of class I myosins expression in gdT lymphocytes from data available at https://www.immgen.org/. **B)** Myo1e and Myo1f expression in total IEL by western blot. **C)** Densitometric analysis of Myo1e and Myo1f expression in total IEL, actin was used as a loading control. **D)** Representative histogram of Myo1e and Myo1f expression in total and γδT IEL by flow cytometry. **E)** Mean Fluorescence Intensity (MFI) of Myo1e and Myo1f expression by flow cytometry. Heat map generated with Prism version 8. The western blot represents one assay with three different mice. Flow cytometry data are from three independent experiments. A t-Student test was applied, p-value; *=0.0142, ***=0.001, ****=0.0001.

### Myo1f deficiency has an impact on small intestinal intraepithelial lymphocyte counts

IELs were obtained from Myo1e−/−, Myo1f−/−, and dKO mice and were stained for flow cytometry analysis to gain clues about the functions of long-tailed class I myosins. First, the t-SNE (t-Stochastic Neighborhood Embedding) algorithm shows that Myo1e did not considerably affect the proportions of IELs. However, Myo1f deficiency negatively affected CD8αα γδT, CD8αβ, and CD8αα αβT IELs while positively affecting CD4 αβT cells and ILCs. In contrast, the absence of both myosins affected CD8 negatively and CD8αβ αβT and positively ILCs IELs. Even though CD8αα γδT IEL proportion was affected in Myo1f absence, the lack of both myosin had no consequence in the proportion of this population (Fig. 2A). To confirm previous data, conventional gate strategy analysis, and IELs counts were performed (Fig. S1). Total IELs were reduced in Myo1e−/−, Myo1f−/− and dKO mice (Fig. 2B). This reduction was also observed for total, CD8αα, and CD8αβ γδT IELs as well as CD8αα, CD8αβ, and CD4 αβT cells (Figs. 2C, S2B). Moreover, the reduction was more prominent in Myo1f than Myo1e, but dKO mice do not have an additive effect. In agreement with the bioinformatics and t-SNE analysis, this suggests that Myo1f has a more relevant role in IEL functions. Fluorescence staining of the small intestine tissue sections confirmed a reduction in TCRγδ+ lymphocytes in the epithelial cell layer and lamina propria in Myo1f−/− mice (Figs. 2D, E). Finally, variable-specific flow cytometry analysis of γδT IEL showed that Myo1f deficiency affects Vγ1, Vγ4, and Vγ7 γδT IELs; however, the reduction was more prominent in the Vγ7 subset in agreement with IELs repertory and Myo1f expression (Figs. 2F, 1A, S1C). Thereby, Myo1f deficiency affects IEL numbers in the small intestine.

**Figure 2.**
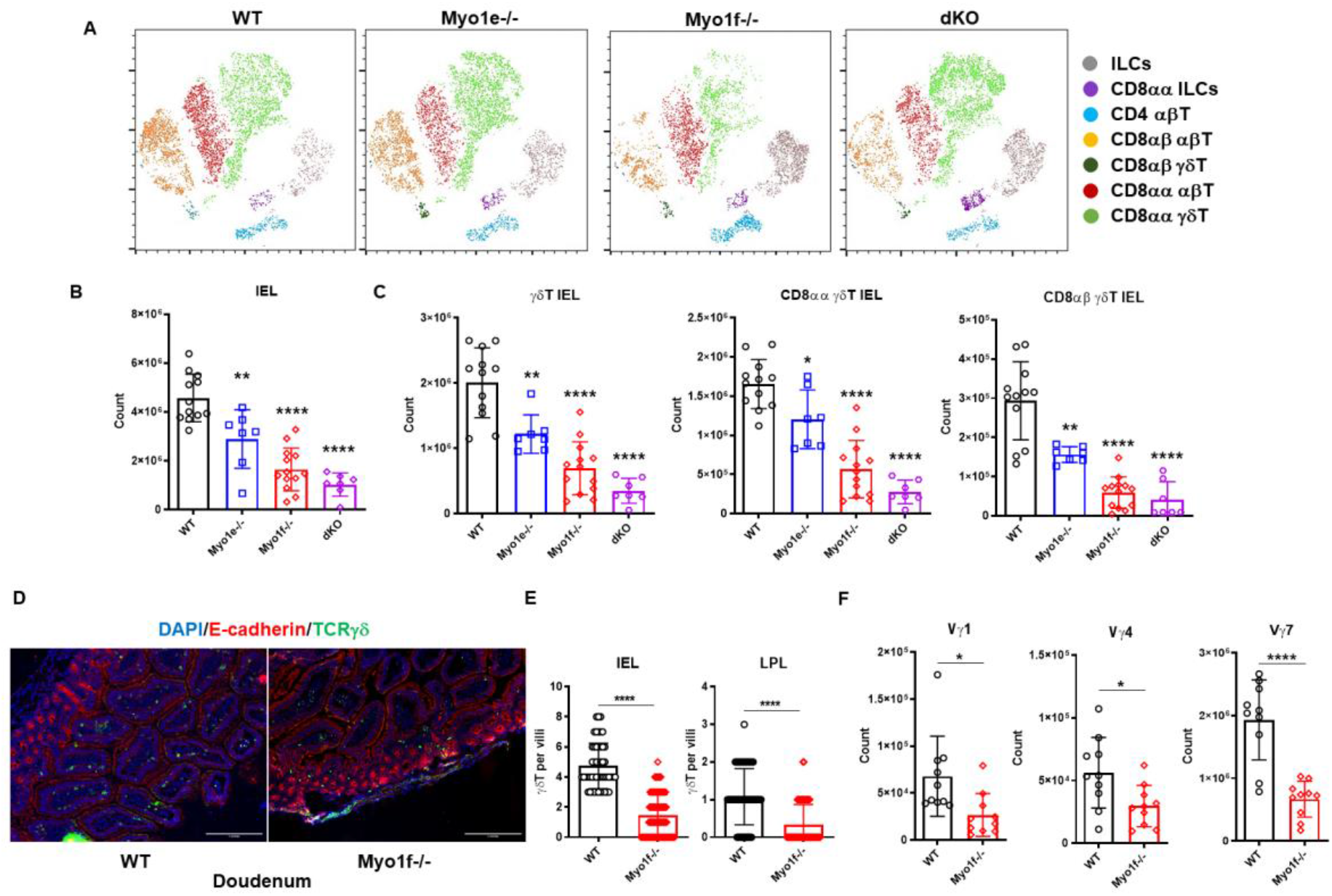
Intraepithelial lymphocyte count in WT, Myo1e−/−, Myo1f−/−, and dKO mice. **A)** t-SNE algorithm analysis from flow cytometry data in WT, Myo1e−/−, Myo1f−/−, and dKO mice. **B)** Total and **C)** γδT, CD8αα γδT and CD8αβ γδT IEL count in WT, Myo1e−/−, Myo1f−/− and dKO mice. **D)** Representative histofluorescense staining of γδT lymphocytes in duodenum from WT and Myo1f−/−. **E)** γδT IEL and LPL count in histofluorescense sections. **F)** Vγ-specific γδT IEL subsets in WT and Myo1f−/− mice. In flow cytometry analysis, each dot represents one mouse. In tissue count, each dot represents villi and shows pooled data from 3 independent experiments. A t-Student test was applied, p-value; *=0.05, **=0.005, ***=0.0005, ****=0.0001

### Myo1f absence affects gut homing receptors and integrin expression

The homing and further localization of IELs in the small intestine is CCR9 and α4β7 integrin-dependent. Therefore, CCR9 and α4β7 integrin expression was evaluated in thymic and γδT IELs from WT and Myo1f−/− mice to identify possible defects in gut homing receptors. Although thymic γδT lymphocytes do not have CCR9 and α4β7 expression changes, γδT IELs positive to CCR9 and α4β7 were reduced in Myo1f−/− (Figs. 3A, B, S4). Furthermore, because Myo1f has been associated with integrin expression, we also evaluated the αE, αLβ2, β1, and αM expression (Fig. 3C). Only a reduction in αLβ2+ γδT IELs in Myo1f−/−was found. In contrast, only the membrane level of αE was slightly affected (Figs. 3D, E). Hence, Myo1f is required for right gut-homing receptor and integrins expression.

**Figure 3.**
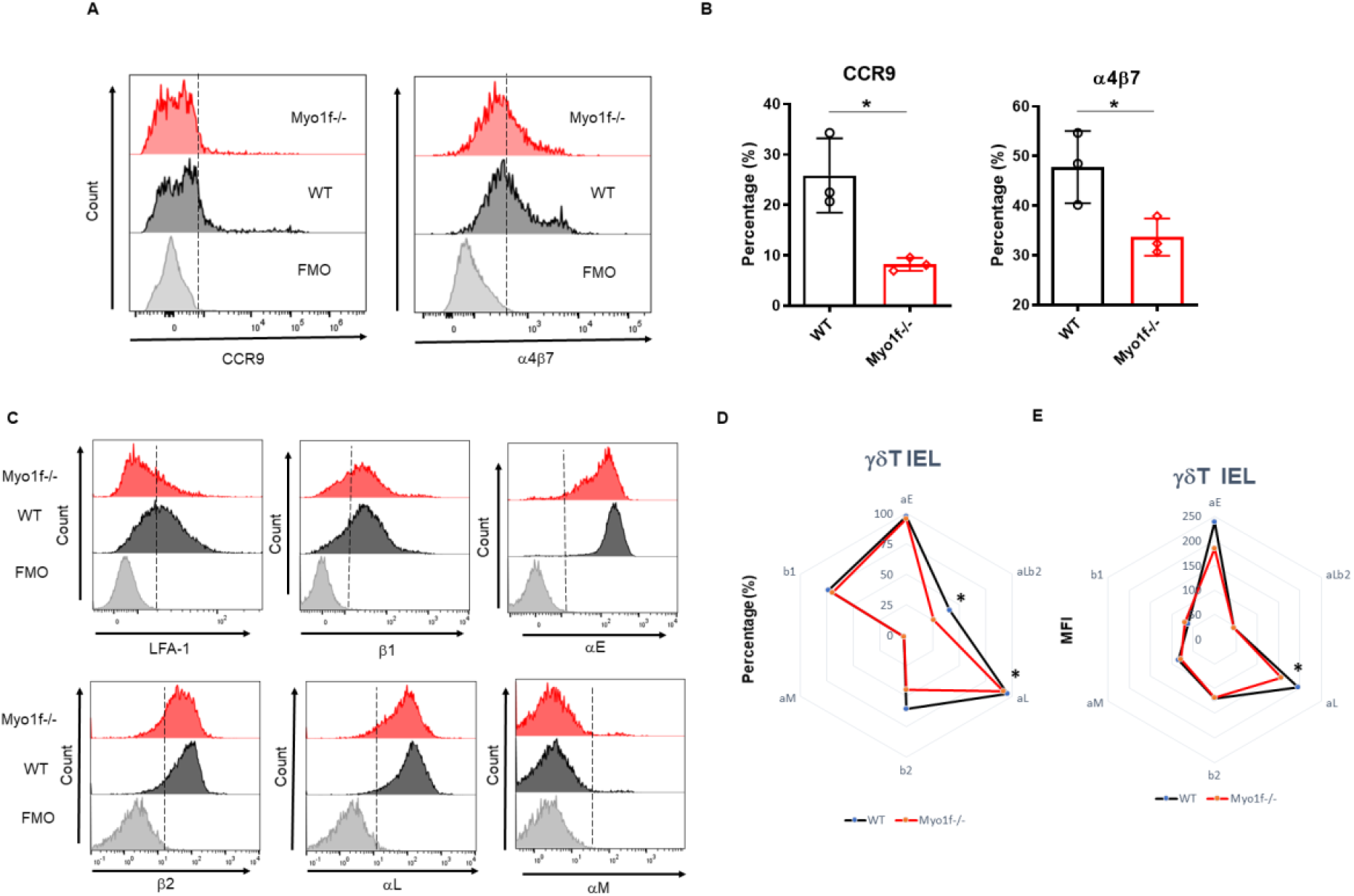
Gut homing and integrin expression. **A)** Representative histogram of surface CCR9 and α4β7 integrin expression in WT vs. Myo1f−/− γδT cells. **B)** Percentage of CCR9 and α4β7 positive cells. **C)** Representative histograms of αLβ2, αL, β2, αM, and β1 integrin expression. **D)** Radar plot of the proportion of γδT cell integrin positive cells. **E)** Radar plot of MFI of integrin+ γδ T cells. Each dot represents one mouse. In radar plots, each plot represents the mean value. WT vs. Myo1f−/− comparison was applied by the t-Student test.

### Myo1f modulates adhesion, spreading, and filopodia formation

Integrins are relevant in cell adhesion. Cell adhesion to recombinant MadCAM-1Fc, fibronectin, and collagen was analyzed to evaluate the consequence of the integrin expression reduction. In agreement with reduced integrin expression, cell adhesion to MadCAM-1, fibronectin, and collagen was reduced in Myo1f deficient γδT cells (Fig. 4A), showing defects in integrin-mediated adhesion. Cytoskeleton polymerization was analyzed by structured illumination microscopy (SIM) to gain clues about Myo1f role in integrin-mediated adhesion. Myo1f−/− γδT cells showed less actin polymerization, and fewer membrane projections (Fig. 4B). MFI of phalloidin staining confirmed less actin polymerization in Myo1f−/− (Fig. 4C). Morphologic analysis showed no defects in area and shape. However, a reduced cellular perimeter was observed (Figs. 4D, E). Reduced perimeter results from shorter filopodia length formation in Myo1f deficient γδT cells (Fig. 4F). These results suggest that Myo1f modulated cell adhesion, spreading, and filopodia formation.

**Figure 4.**
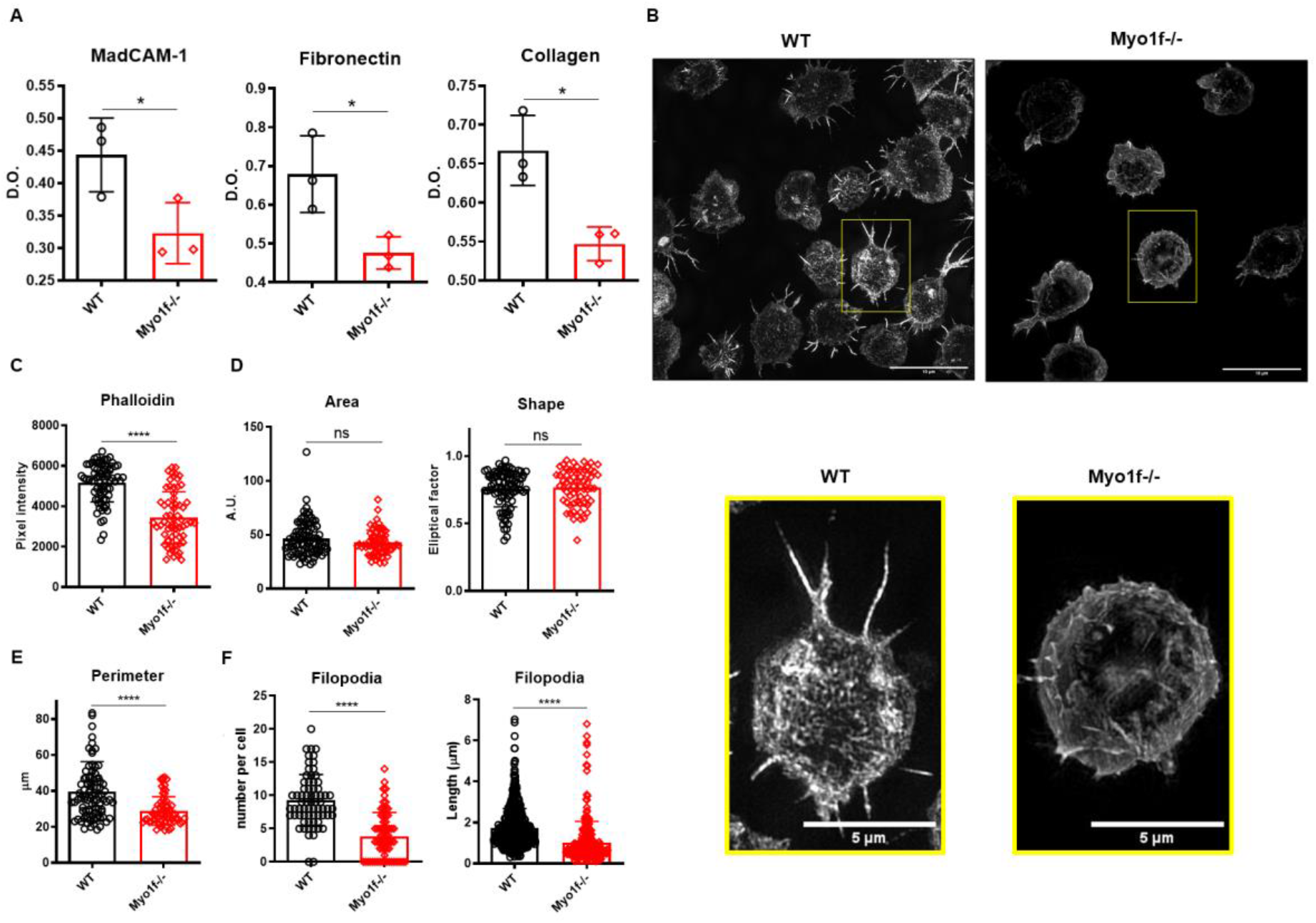
Cell adhesion and spreading. **A)** Cell adhesion to MadCAM-1, fibronectin, and collagen I. **B)** Representative images of phalloidin staining by structured illumination microscopy (SIM). **C)** Phalloidin pixel intensity. **D)** Cell morphologic analysis of the area, shape, and **E)** Perimeter. **F)** Filopodia count per cell and length. In cell adhesion, each dot represents an average of 2 duplicates in 3 independent experiments. In morphologic analysis, each dot represents one cell polled from 3 independent experiments. A t-Student was applied, p-value; *=0.05, **=0.005, ***=0.0005, ****=0.0001.

### Myo1f has a role in migration and lamellipodia formation

To determine if Myo1f has a role in γδT cell migration, we evaluated 2D CCL25-independent and -dependent migration. Myo1f−/− cells showed reduced migration in the absence of CCL25 (Fig. 5A), denoted by a reduced accumulated (total distance traveled between two points) and Euclidian (straight line distance between two points) distance, and velocity (Fig. 5B). CCL25-dependent migration showed a gradient-specific migration by WT cells. The CCL25-dependent migration by WT cells was reduced with an anti-CCR9-blocking antibody. Myo1f-deficient cells showed reduced migration to the CCL25 gradient, consistently with a reduced expression of CCR9, and comparable to CCR9-blocked WT cells (Fig. 5C).

**Figure 5.**
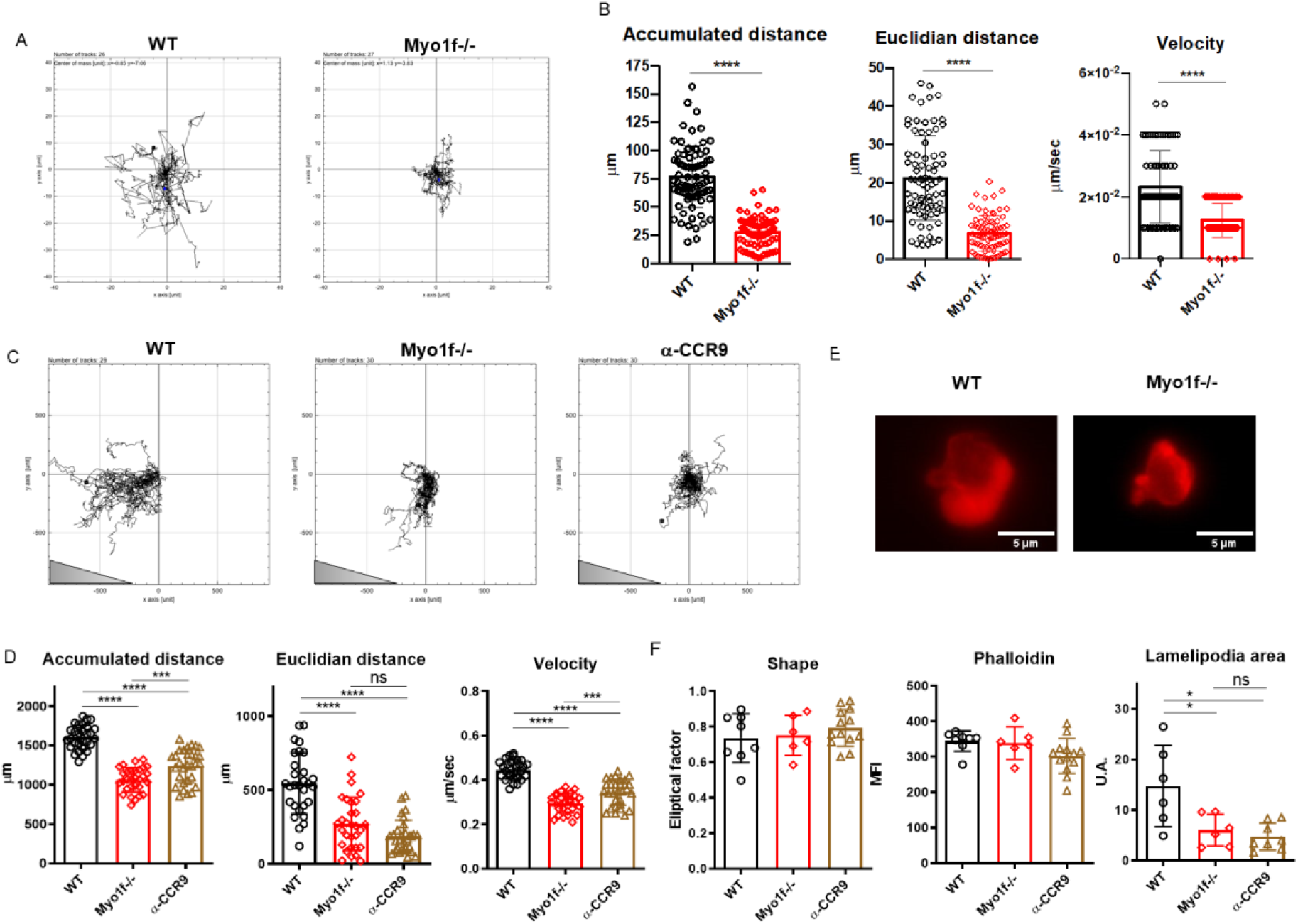
CCL25-independent and -dependent migration. **A)** Representative single-cell migration tracks in CCL25-independent migration. **B)** Accumulated and Euclidian distance and velocity. **C)** Representative single-cell migration tracks in CCL25 dependent migration. **D)** Accumulated and Euclidian distance and velocity in CCL25-dependent migration. **E)** Phalloidin staining of γδT lymphocyte migratory phenotype during CCL25-dependent migration. **F)** Phalloidin MFI, shape factor, and lamellipodia area of γδT lymphocyte migratory phenotype. Each dot represents one cell, a pool of 3 independent experiments. Triangle represents gradient orientation. A t-Student test was applied, p-value; *=0.05, **=0.005, ***=0.0005, ****=0.0001.

Myo1f−/− γδT cells demonstrated reduced accumulated and Euclidian distance and velocity (Fig. 5D). Myo1f−/− γδT cells showed a similar shape and total actin polymerization. However, they had a reduced lamellipodia area compared with WT γδT cells (Figs. 5E, F). Thus, Myo1f is required in chemokine-independent and - dependent migration and lamellipodia formation.

### Myo1f regulates CCR9 and α4β7 polarization and signaling

To elucidate the defects in the mechanism of adhesion and migration in Myo1f−/− γδT IEL, we evaluated the CCR9 and α4β7 integrin polarization by capping assays. WT γδT IEL polarizes CCR9 receptor and α4β7 integrin in large patches, whereas Myo1f−/− γδT IEL showed smaller scattered patches (Fig. 6A).

**Figure 6.**
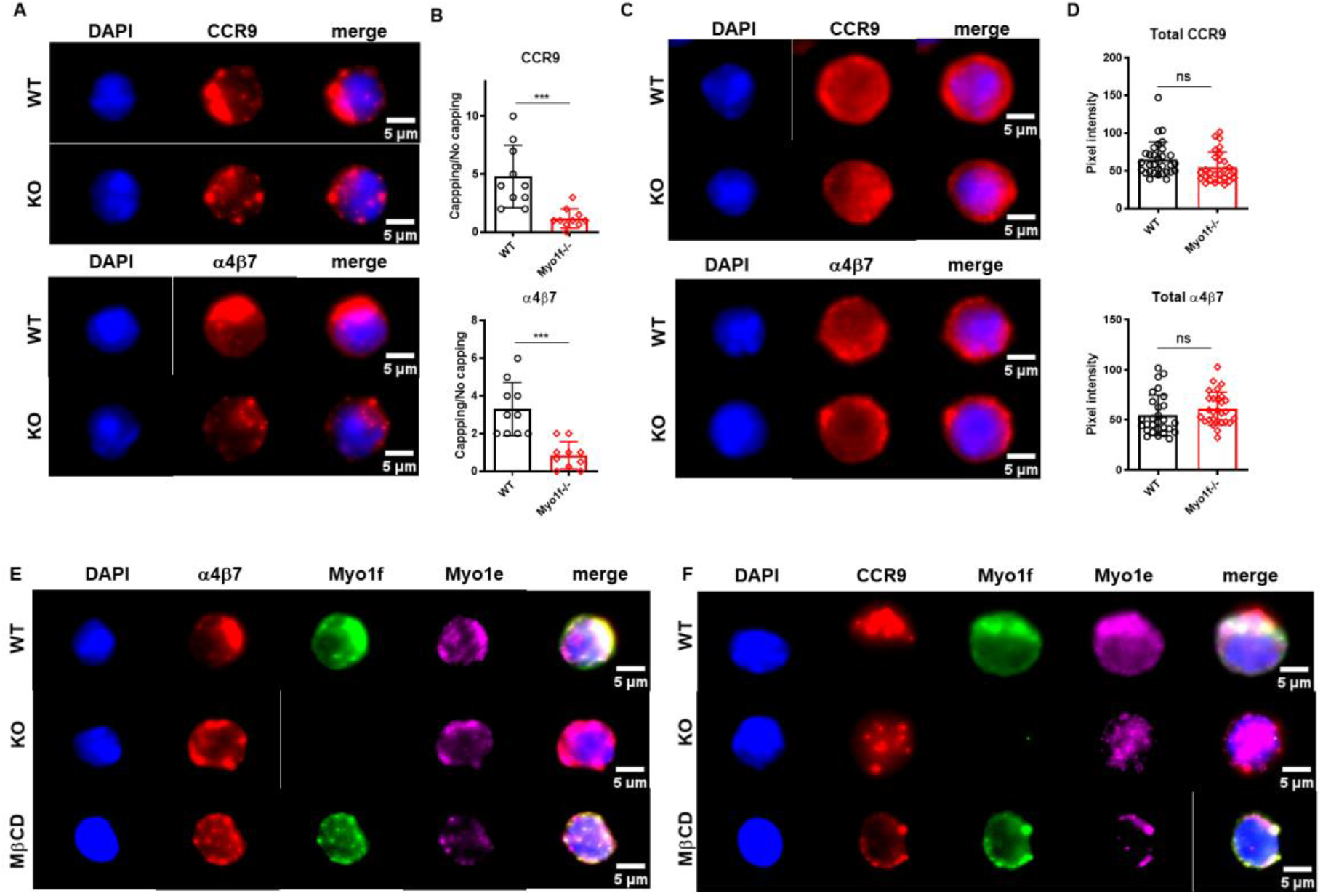
Myo1f in CCR9 and α4β7 integrin polarization. **A)** Representative CCR9 and α4β7 capping induction in WT and Myo1f−/− γδT IEL. **B)** Capping/no capping quotient of CCR9 and α4β7 **C)** Total CCR9 and α4β7 staining in WT and Myo1f−/− γδT IEL **D)** Pixel intensity of total CCR9 and α4β7 staining **E)** Representative α4β7 capping phenotypes from WT, Myo1f−/− and MβCD pre-treated γδT IEL. **F)** Representative CCR9 capping phenotypes from WT, Myo1f−/−, and MβCD pre-treated γδT IEL. Poled data from 3 independent experiments. A t-Student test was applied, p-value; ***=0.0005.

The ratio of cells with capping/uncapping images confirms that Myo1f deficient cells fail to polarize CCR9 and α4β7 (Fig. 6B). Surface expression of CCR9 and α4β7 was reduced in Myo1f defective cells compared to WT concordantly with flow cytometry data (Fig. S4A, B, 3A, B). In contrast, total staining remains identical, suggesting no defects in the de novo synthesis of both molecules (Fig. 6C, D).

Previously, Myo1g has shown accumulation in clustering sites of adhesion molecules (López-Ortega and Santos-Argumedo, 2017). To investigate where long-tailed class I myosin localize during the capping process Myo1e and Myo1f staining was performed. Myo1e and Myo1f localize in CCR9 and α4β7 in WT γδT IEL, whereas Myo1e remains adjacent to the capping site in α4β7 but has a diffuse localization in the CCR9 capping site in Myo1f−/− γδT IEL (Fig. 6E, F). This result suggests that Myo1e and Myo1f participate in protein polarization at the plasma membrane. Protein clustering in the cell membrane requires cholesterol and sphingolipid-enriched liquid-ordered microdomains called lipid rafts (Simons and Roomre, 2000). To evaluate the role of lipids rafts, WT γδT IEL were preincubated with methyl-β-cyclodextrin (MβCD) before capping assay. MβCD phenotype resembles Myo1f−/− smaller and scattered patches of CCR9 and α4β7 (Fig. 6E, F). In contrast, both myosins remain localized at the capping site at CCR9 and α4β7 (Fig. 6E, F). This result suggests that long-tailed class I localization at the capping site does not depend on the lipid raft ensemble.

CCR9 and α4β7 clustering, as well as lipid raft, are essential in signal transduction (Simons and Roomre, 2000). Therefore, tyrosine phosphorylation induced by CCR9 and α4β7 was evaluated to determine the role of Myo1f in signaling. As expected, CCR9 and α4β7 deficient-polarization clustering result in a reduced phosphorylation cascade. Similarly, MβCD phenotype resembles Myo1f−/− reduced tyrosine phosphorylation (Fig. 7A, B), denoted by less phosphorylation area and fluorescence intensity (Fig. 7C, D). Thus, Myo1f regulates CCR9 and a4b7clustering in the plasma membrane to induce phosphotyrosine-mediated signal transduction.

**Figure 7.**
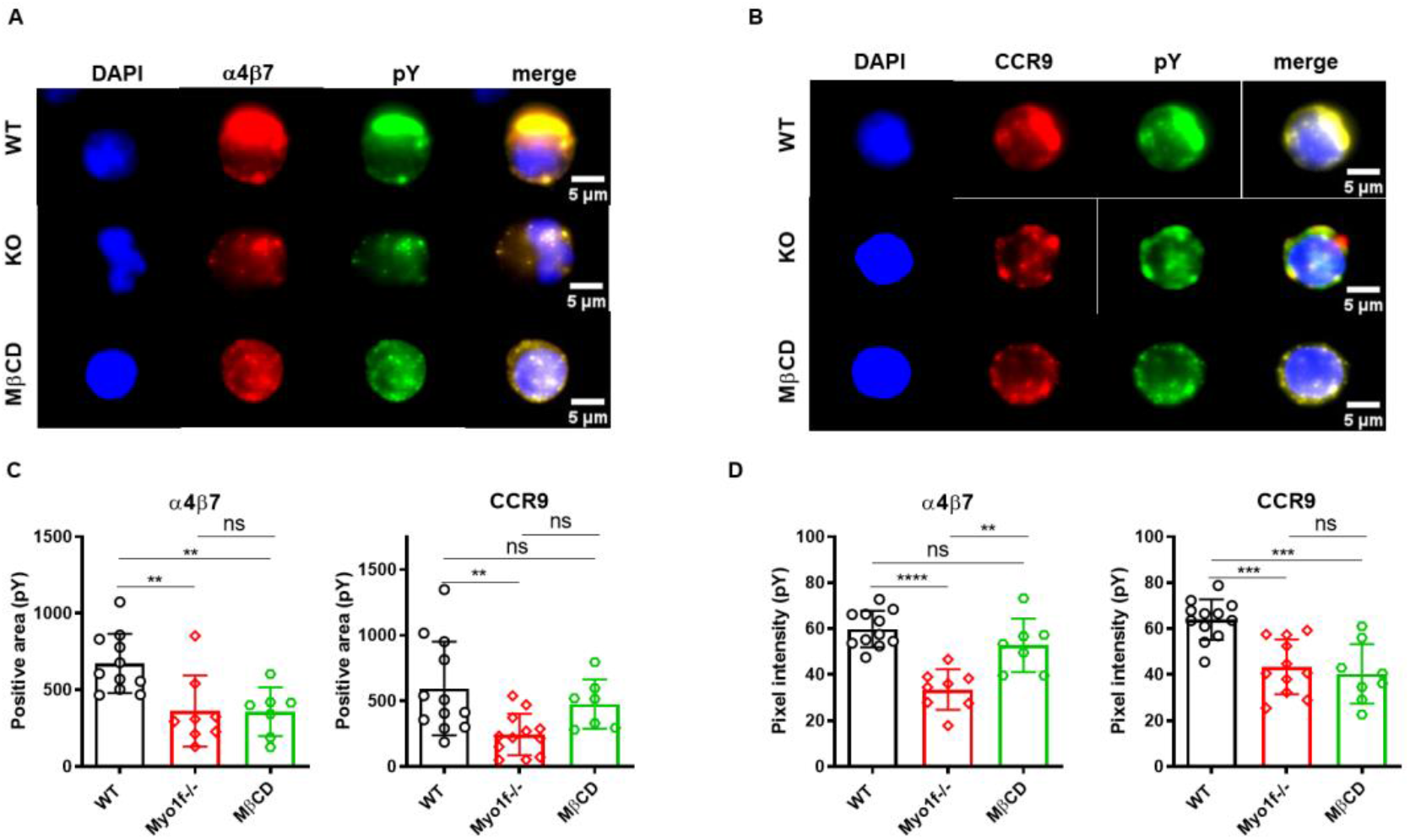
Myo1f deficiency in tyrosine phosphorylation-mediated signaling. **A)** Representative α4β7 capping phenotype and phosphotyrosine staining from WT, Myo1f−/− and MβCD pre-treated γδT IEL **B)** Representative CCR9 capping phenotype and phosphotyrosine staining from WT, Myo1f−/− and MβCD pre-treated γδT IEL **C)** Phosphotyrosine positive area in WT, Myo1f−/− and MβCD pre-treated γδT IEL in α4β7 and CCR9 capping phenotype **D)** Phosphotyrosine pixel intensity in WT, Myo1f−/− and MβCD pre-treated γδT IEL in α4β7 and CCR9 capping phenotype. Poled data from 3 independent experiments. One-way ANOVA test was applied, p-value; **=0.005, ***=0.0005, ****=0.0001.

## DISCUSSION

Class I myosins have been related to migration, adhesion, and vesicular trafficking in leukocytes (Girón-Pérez et al., 2019; Navinés-Ferrer and Martín, 2020). However, their role in T lymphocytes remains poorly addressed. This work addresses the role of long-tailed class I myosins in the adhesion and migration of intraepithelial lymphocytes.

Here, we demonstrated that Myo1e and Myo1f are expressed at the protein level by αβT and γδT intraepithelial lymphocytes. Myo1f is more abundant than Myo1e in these cells. Consistently with the data available in the Immunological Genome Project (https://www.immgen.org/), neither thymic nor spleen γδT cells express Myo1e, but ∼25% of cells express Myo1f. Additionally, activated γδT cells increased Myo1f expression compared to their naïve counterpart. These suggest that long-tailed class I myosins could be expressed after activation or differentiation. It has been previously demonstrated that monocytes acquire Myo1f expression upon macrophage differentiation (Piedra-Quintero et al., 2019). At the same time, Myo1e deficiency defects in B cells are activation-dependent (Girón-Pérez et al., 2020). In this regard, inverse Myo1f and Tasp1 relative expression correlate with differentiation in stem cell-monocyte-macrophage lineage. Mechanistically, Myo1f is cleaved by the caspase 1 protease, which impaired its function, indicating a Tasp1 negative regulation of Myo1f during differentiation (Hensel A. et al., 2022). As effector cells, γδT intraepithelial lymphocytes express more Myo1e and Myo1f than their thymus and spleen counterpart.

Myo1e−/− and Myo1f−/− mice have reduced intraepithelial lymphocytes in the small intestine. However, only Myo1f−/− mice have a decreased γδT lymphocyte proportion, whereas Myo1e−/− mice remain identical to WT. This difference could be due to Myo1f showing higher expression than Myo1e. Additionally, Myo1e and Myo1f are different in the 950-1000 amino acid region, where the TH2 domain is located, and thus, this could imply different protein-protein interactions (https://www.uniprot.org/). However, this needs to be experimentally confirmed. Despite this, dKO mice have no additive effects in the γδT cell proportion. It has been demonstrated that dKO macrophages showed increased actin waves and Arp2/3 recruitment during Fc-mediated phagocytosis (Barger et al., 2019), suggesting altered actin polymerization. If the long-tailed class I myosin deficiency is compensated by other class I myosins requires to be addressed. Based on these results, we focused on understanding the Myo1f function in γδT intraepithelial lymphocytes.

The α4β7 integrin is essential in gut-specific recruitment (Wagner et al., 1996). B7−/− mice have neither intraepithelial lymphocytes, abT, or γδT in the gut (He et al., 2019). Myo1f−/− γδT intraepithelial lymphocytes express less α4β7 integrin. Similarly, the knock-down of Myo1f in mast cells reduces β7 integrin expression (Navinés-Ferrer et al., 2019). Contrary to the first description of Myo1f deficiency in neutrophils (Kim et al., 2006), αLβ2 integrin was decreased in Myo1f−/− γδT intraepithelial lymphocytes. This difference could be explained due to the cell type or activation status. Moreover, β2−/− mice have no defects in γδT or αβT intraepithelial lymphocyte count (McIntyre et al., 2020). We did not observe significant differences in β1 and αM expression. However, β1 low expression was reported in Myo1f knock-down mast cells (Navinés-Ferrer et al., 2019), while αVβ3 integrin expression is enhanced in stably-transfected macrophages with Myo1f-GFP (Piedra-Quintero et al., 2019). Our results provide evidence of the relevant role of Myo1f in integrin expression.

Due to decreased integrin expression, Myo1f−/− γδT IEL showed less adhesion to MadCAM-1 (ligand of α4β7), fibronectin, and collagen. Mechanically, Myo1f−/− deficiency results in fewer filopodia formation. This filopodia formation depends mainly on the actin cytoskeleton polymerization dependent on the activity of Cdc42 (Sit and Manser, 2011). In agreement, reduced Cdc42 activation was found in resting and IgE-mediated exocytosis (Navinés-Ferrer et al., 2021).

Intraepithelial lymphocyte homing depends on CCR9, and its deficiency reduces γδT proportion (Wurbel et al., 2001; Uehara et al., 2002). Similarly, CCL25 deficiency showed a similar reduction (Wurbel et al., 2007). We demonstrated that Myo1f−/− deficiency results in less CCR9 expression. 2D migration assays showed less accumulated and Euclidian distance and velocity in a CCL25-dependent and independent migration. These results suggest that Myo1f is required for chemokine-dependent and independent migration. Myo1f−/− cells have reduced lamellipodia area in the CCL25-dependent migration, possibly due to a reduction in the Rac1 activation. Rac1 regulates lamellipodia formation by actin polymerization during migration (Sit and Manser, 2011). Myo1e−/− B cells stimulated with CXCL12 show reduced Rac1 activation (Girón-Pérez et al., 2020). However, the precise mechanism by which Myo1f-dependent Rho GTPases activation occurs is unknown. Myo1f interacts with 3BP2 (Navinés-Ferrer et al., 2019), an adapter molecule necessary in neutrophil migration (Chen et al., 2012) which is essential in Rho GTPase family activation by modulating VAV1 activation (Foucault et al., 2005), suggesting that Rho GTPase activation defects in Myo1f deficiency could be due to VAV reduced activation. This way, long-tailed class I myosins are relevant for the Rho GTPase family activation.

Typically, class I myosin localize in the plasma membrane and colocalizes with cortical actin. This localization has been related to membrane tension regulatory properties. In this regard, it has been reported that Myo1g controls plasma membrane tension in B cells (López-Ortega et al., 2016). Additionally, Myo1g participates in lipid raft-CD44 mobilization and recycling (López-Ortega and Santos-Argumedo, 2017). Here we demonstrate that Myo1f deficiency impaired correct CCR9 and α4β7 polarization. Moreover, Myo1e and Myo1f relocalize at the capping site, indicating their participation in this process. Furthermore, cholesterol depletion with MβCD in WT cells resembles Myo1f−/− capping phenotype suggesting lipid raft-mediated polarization. Despite impaired CCR9 and α4β7 polarization in MβDC preincubation, Myo1f localization remains at the capping site, suggesting that Myo1f localization is lipid raft-independent. However, MβCD disrupts Myo1e localization at the CCR9 capping site. The specific phosphoinositide binding by Class I myosins could explain the differences in the patterns observed between Myo1e, and Myo1f in γδT intraepithelial lymphocytes (Chen, C.L. et al., 2012).

Finally, we demonstrated that altered CCR9 and α4β7 polarization reduce tyrosine phosphorylation compared to the WT counterpart. Together, this work provides evidence about the role of Myo1f in adhesion and migration in γδT intraepithelial lymphocytes, mediating the lipid raft dependent-CCR9 and α4β7 polarization and cell signaling.

## MATERIAL AND METHODS

### Mice

Female (6-8 weeks old) C57BL/6 wild type, Myo1e−/− (B6.129S6(Cg)-*Myo1e*^*tm1*.*1Flv*^/J), Myo1f−/− (B6.129S6-*Myo1f*^*tm1*.*1Flv*^/J) and Myo1e−/−+Myo1f−/− (dKO) (generated in the Cinvestav by crossing Myo1e−/− with Myo1f−/− during ten generations to obtain double homozygous knock out) mice were bred and maintained at Cinvestav-UPEAL (Unit of Production and Experimentation with Animals of Laboratory) (Mexico City, Mexico) facilities. Myo1f−/− were maintained in the Johns Hopkins Bloomberg School of Public Health (Baltimore, Maryland, USA) facilities for some experiments. WT mice were purchased from Jackson Laboratories. Myo1e−/− and Myo1f−/− mice were kindly donated by Dr. Richard Flavell from Yale School of Medicine, New Haven, CT. Mice genotypification was carried out employing the primers described previously (Kim et al. 2006; Krendel et al., 2009).

All experiments were approved by the Animal Care and Use Committee at Cinvestav and carried out following ARRIVE (Animal Research: Reporting of *in vivo* experiment) guide.

### Bioinformatic Analysis

Class I myosin expression by γδT lymphocytes was obtained from The Immunological Genome Project from the portal https://www.immgen.org. mRNA expression was manually obtained and graphed as a heat map using Prism Software (Irvine, CA) version 8.

### Intraepithelial lymphocyte isolation

Complete small intestines were removed from mice, cut longitudinally, and then in 1 cm fragments. Then, fragments were placed in 1 mM DTE (1,4-Dithioerythritol) (Sigma-Aldrich, St Louis, MO.) in HBSS (Hanks Balance Salt Solution) without Ca^+2^ and Mg^+2^ (Thermo Fisher Scientific, Waltham, MA.) for 30 min, twice. The cell suspension was filtered using a 70 μM cell strainer (Thermo Fisher Scientific, Waltham, MA.), transferred to 50 mL tubes, and centrifuged at 1500 rpm for 10 min. Pellets were washed twice with HBSS and then resuspended in discontinuous 40/70% Percoll® (Sigma-Aldrich, St Louis, MO.) gradient and centrifuged at 2500 rpm for 30 min (Qui and Sheridan, 2018). The interface was collected and washed two times.

### Flow Cytometry

IELs were incubated with anti-CD16/32 (clone 93, dilution 1:500) antibody (Biolegend, San Diego, CA.) and Live/Dead Aqua (dilution 1:1000) (Thermo Fisher Scientific, Waltham, MA.) for 30 min on ice. Then, IELs were washed and incubated with anti-CD45 (clone 30-F11, dilution 1:500) (eBioscience, Thermo Fisher Scientific, Waltham, MA.), - CD3ε (clone 145-2C11, dilution 1:300) (eBioscience), - TCRγδ (clone GL3, dilution 1:300) (BD Biosciences, Franklin Lakes, NJ.), - CD8α (clone 53-6.7, dilution 1:300) (BD Biosciences), - CD8β (clone YTS156.7.7, dilution 1:300) (Biolegend) and CD4 (clone GK1.5, dilution 1:200) (BD Biosciences) for total IEL analysis. With anti-CD3ε (eBioscience), - TCRγδ (BD Bioscience), - TCRVγ1.1/1.2 (clone 4B2.9, dilution 1:200) (Biolegend), - TCRVγ2 (clone UC3-10A6, dilution 1:200) (BD Biosciences) for Vγ specific γδT IEL analysis. Anti-CCR9 (clone eBioCW-1.2, dilution 1:200) (eBioscience), - α4β7 (clone DATK-32, dilution 1:200) (Invitrogen, Thermo Fisher Scientific, Waltham, MA.), - αE (clone 2E7, dilution 1:200) (eBioscience), - αLβ2 (clone H155-78, dilution 1:200) (Biolegend), - αM (clone M1/70, dilution 1:200) (Biolegend), - β1 (clone HMb1-1, 1:200) (Biolegend), were also employed. For intracellular staining, cells were fixed with 1% paraformaldehyde (PFA) for 5 min. Then the cells were permeabilized with saponin 0.1% for 15 min, next incubated with 10% goat serum (Sigma-Aldrich, St. Louis, MO.) at 37 °C, and further incubated with anti-Myo1e (Cat. PAD434Mu01, dilution 1:500, Cloud-Clone Corp, Houston, AZ.), - Myo1f (Cat. 13933-AP, dilution 1:500) Proteintech, Rosemont, IL.). For indirect staining, secondary anti-rabbit FITC (dilution 1:1000, Jackson ImmunoResearch Laboratories, West Grove, PA.) or anti-rabbit Pacific Blue (dilution 1:1000 Jackson ImmunoResearch Laboratories) were employed. Finally, cells were resuspended in PBS and analyzed in Cytoflex LX (Beckman Coulter, Brea, CA.), LSR Fortessa (Becton Dickinson, San Jose, CA.), BD Influx (Becton Dickinson, San Jose, CA.) or Attune Next (Thermo Fisher Scientific, Waltham, MA.) cytometers. Data were analyzed using FlowJo Software (BD Biosciences) version 10.6.

### γδT IEL purification and culture

IELs were incubated with anti CD16/32 for 30 min at 4 °C. Then, cells were incubated with anti-TCRαβ biotin-coupled (H57-597) (Biolegend) for 30 min at 4 °C. Next, cells were incubated with anti-biotin beads (Miltenyi Biotec, Bergisch Gladbach) for 15 min at 4 °C. Finally, cells were washed, and γδT IELs were negatively selected by an LD MACS column (Miltenyi Biotec). γδT IELs were incubated with Live/Dead Scarlet (Thermo Fisher Scientific) and anti-TCRγδ-PE (Biolegend) antibody for 30 min at 4 °C. Cell viability and purity were confirmed by flow cytometry. Cell viability was >95%, and purity was 90%. IELs were cultured as described previously (Swamy M. et al., 2015). Purified γδT IELs were incubated with RPMI 1640 (Thermo Fisher Scientific) medium supplemented with 10% fetal bovine serum (ATCC, Manassas, VA.), 2.5% HEPES (Thermo Fisher Scientific), 1% glutamine (Thermo Fisher Scientific), 1% Pen/Strep (Thermo Fisher Scientific), 1% sodium pyruvate (Thermo Fisher Scientific), 1% non-essential amino acids (Thermo Fisher Scientific) and 0.2% β-mercaptoethanol (Thermo Fisher Scientific) plus murine recombinant IL-2 (10 U/mL), IL-4 (200 U/mL), IL-3 (100 U/mL), and IL-15 (100 U/mL) (PeproTech, Cranbury, NJ.). After two days, cells were re-plated and cultured only with IL-2 (10 U/mL). The medium was replaced every 3-5 days.

### Western blot

Proteins from total extracts (30 μg) were resolved in 10% SDS-PAGE gels. Proteins were transferred to a nitrocellulose membrane (Bio-Rad, Hercules CA.), and protein transfer was confirmed with Ponceau red. First, membranes were blocked with 5% milk in 0.05 % TBS-Tween 20 buffer for 1 h at room temperature. Next, anti-Myo1e (dilution 1:1000, Cloud-Clone Corp) and -Myo1f (dilution 1:2500, Proteintech) antibodies were incubated in 2% milk in TBS-T overnight at 4 °C; next, membranes were washed and incubated with anti-rabbit HRP (dilution 1:10000, Jackson ImmunoResearch Laboratories) antibodies for 1 h at room temperature. Finally, membranes were washed and revealed with a super signal western blot system (Pierce™, Thermo Fisher Scientific, Waltham, MA.) in C-digit equipment (LI-COR Bioscience, Cambridge, UK.). Densitometric analysis was performed using Image J software (National Institutes of Health, Bethesda, MD.) version 1.52.

### Tissue immunofluorescence staining

Duodenum sections were obtained from WT and Myo1f−/− mice. First, tissue fractions were pulled in optimal cutting temperature media (ProSciTech, Townsville, Queensland, Australia) and frozen in liquid nitrogen. Next, tissue sections (5 μm thick) were placed in poly-L-lysine coated slides, fixed with absolute acetone for 20 min at −20 °C, and then blocked with 5% BSA for 1 h at room temperature. Next, tissues were incubated with purified anti-TCRγδ (dilution 1:500, Biolegend) and anti-E-cadherin (dilution 1:100, Santa Cruz Biotechnology, Dallas, TX.) antibodies for 1 h a room temperature. Afterward, tissues were stained with anti-hamster-AF555 (dilution 1:1000, Thermo Fisher Scientific) and anti-rat-FITC (dilution 1:1000, Jackson ImmunoResearch Laboratories) secondary antibodies, respectively. Finally, tissues were incubated with DAPI (dilution 1:10000, Thermo Fisher Scientific) for 10 min and mounted with 10% glycerol. Tissues were analyzed immediately in an Olympus X microscope (Olympus Scientific, Tokyo, Japan). γδT cell count was performed with Image J software (NIH), version 1.52.

### Cell adhesion Assay

Plates (96-wells) were pre-coated with murine recombinant MadCAM-1 (R&D Systems, Minneapolis, MN.), fibronectin, or collagen overnight at 4 °C. Then, plates were blocked with 5% BSA in PBS for 1 h a 37 °C. Next, 2 × 10^5^ γδT IELs were seeded and incubated at 37 °C for 1 h. After that, non-adhered cells were removed, and plates were washed once with PBS. Then cells were fixed with PFA 4% for 10 min at room temperature. Next, plates were washed and stained with crystal violet for 30 min; plates were washed 5 times and then incubated with 10% SDS for 30 minutes. Finally, plates were read in a 680 microplate reader (Bio-Rad, Hercules CA.) reader plate equipment.

### Spreading

γδT IELs (1 × 10^5^) were seeded in 1.5 mm coverslips previously coated with 5 μg/mL of Collagen I (Corning®, Bedford, MA.). Cells were incubated for 1 h at 37 °C with 5% CO2. Next, cells were fixed with 4% formaldehyde for 30 minutes at room temperature and then permeabilized with 0.2% Triton X-100 (Sigma-Aldrich) for 15 minutes. After that, cells were stained with phalloidin-CF555® (dilution 1:500, Biotium, San Francisco, CA.) for 1 h at 37 °C and mounted with ProlongGold with DAPI (Thermo Fisher Scientific) Cells were observed in a GE DeltaVision OMX-SR microscope (GE Healthcare, Chicago, IL). Images were analyzed using Fiji Software (National Institutes of Health, Bethesda, MD.) version 2.3.0.

### 2-dimension migration assay

γδT IELs (2 × 10^5^) were seeded in an μ-Slide Chemotaxis® chamber coated with collagen IV (Ibidi, Martinsried, Munich, Germany) for 30 min at 37 °C. Then, non-adhered cells were removed, and the chamber was filled with a complete RPMI-1640 medium. Murine recombinant CCL25 (R&D Systems, Minneapolis, MN.) (100 ng/ml) was added in one slide plug. Slides were immediately placed on the platen of a DeltaVision Elite microscope (GE Healthcare, Chicago, IL). Time-lapse videos were taken every 30 sec for 30 min. Distance and velocity analysis was made with Fiji software (Fiji Software (NIH), version 2.3.0, with the TrackMate plugin (Tinevez et al., 2017) and the Chemotaxis and Migration Tool (Ibidi).

### Capping Assay

γδT IEL were incubated with purified anti-CCR9 (clone 9B1, dilution 1:100) (Biolegend) or anti α4β7(clone DATK-32, dilution 1:100) antibody for 20 min at 4°C. Cells were washed with PBS and incubated with the anti-rat-AF594 (dilution 1:500, Thermo Fisher Scientific) antibody for 30 min at 37 °C to induce cross-linking. Cells were washed and then fixed with 4% PFA. For intracellular staining, cells were permeabilized with 0.1% Triton X-100 for 15 min, then incubated with 5% BSA for 1 h, and then incubated with anti-Myo1e (dilution 1:200, Proteintech), anti-Myo1f (clone B-5, dilution 1:200) (Santa Cruz Biotechnology) and anti-phospho-tyrosine (clone PY20, dilution 1:1000) (Biolegend) antibodies. Then, cells were incubated with anti-mouse FITC (dilution 1:1000, Thermo Fisher Scientific) and anti-rabbit AF647 (dilution 1:1000, Biolegend). After staining, cells were mounted in poly-L-lysine (Sigma-Aldrich) pre-coated coverslips with ProlongGold with DAPI. In some experiments, cells were pre-incubated with 5 mM MβCD (Sigma-Aldrich) for 15 min at 37 °C. For the phosphotyrosine detection, cells were treated with 1 mM vanadate (New England BioLabs, Ipswich, MA.).

### Statistical analysis

Results shown are mean with +/− SD. An unpaired two-tailed Student’s *t*-test or one-way ANOVA was applied to assess statistical significance between groups. The p values and the number of samples or cells (n) used are mentioned in each figure legend.

## ACKNOWLEDGMENTS

We thank Dr. Richard Flavell for the kind donation of Myo1e−/− and Myo1f−/− mice. We appreciate the contribution of Dr. Fidel P. Zavala and Dr. Yevel Flores-Garcia to this work for the opportunity to conduct experiments at the Johns Hopkins Bloomberg School of Public Health. Dr. Mira Krendel for the kind donation of Myo1f−/− mice to establish a new colony at Johns Hopkins University. We acknowledge the Johns Hopkins School of Medicine Microscopy Facility (MicFac), LaToya Roker, and Barbara Smith for their assistance. Carlos Trujillo, Ivonne Hernández, and Hector Romero for their technical support at Cinvestav.

## AUTHOR CONTRIBUTIONS

Conceptualization: IUMV, LSA, PTR; Methodology: IUMV, MESB, CEMR, FHC; Validation: IUMV, LSA, PTR; Formal analysis: IUMV Investigation: IUMV, MESB, CEMR, FHC; Resources: LSA, PTR; Writing (original draft): IUMV; Review and Editing: IUMV, LSA, PTR; Visualization: IUMV, LSA, PTR; Supervision: LSA, PTR; Funding: LSA, PTR.

## DATA AVAILABILITY

All data and material generated in this work were included either as primary or supplementary material.

## REFERENCES

Barger, S. R., Reilly, N. S., Shutova, M. S., Li, Q., Maiuri, P., Heddleston, J. M., Mooseker, M. S., Flavell, R. A., Svitkina T., Oakes, P.W., et al., (2019). Membrane-cytoskeletal crosstalk mediated by myosin-I regulates adhesion turnover during phagocytosis. Nat. Commun. 10, 1–18.

Boll, G., Rudolphi, A., Spieβ, S., & Reimann, J. (1995). Regional specialization of intraepithelial T cells in the murine small and large intestine. Scand. J. Immunol. 41, 103–113.

Bonneville, M., Janeway, C. A., Ito, K., Haser, W., Ishida, I., Nakanishit, N., & Tonegawa, S. (1988). Intestinal intraepithelial lymphocytes are a distinct set of γδ T cells. Nature, 336, 479–481.

Chen, C. L., Wang, Y., Sesaki, H., & Iijima, M. (2012). Myosin I links PIP3 signaling to remodeling of the actin cytoskeleton in chemotaxis. Sci. Signal. 5, ra10–ra10.

Chen, G., Dimitriou, I., Milne, L., Lang, K. S., Lang, P. A., Fine, N., Ohashi P. S., Kubes P., & Rottapel, R. (2012). The 3BP2 adapter protein is required for chemoattractant-mediated neutrophil activation. J. Immunol. 189, 2138–2150.

Cheroutre, H., Lambolez, F., & Mucida, D. (2011). The light and dark sides of intestinal intraepithelial lymphocytes. Nat. Rev. Immunol. 11, 445–456.

Darlington, D., & Rogers, A. W. (1966). Epithelial lymphocytes in the small intestine of the mouse. J. Anat. 100, 813

Edelblum, K. L., Shen, L., Weber, C. R., Marchiando, A. M., Clay, B. S., Wang, Y., Prinz I, Malissen B., Sperling A.I. & Turner, J. R. (2012). Dynamic migration of γδ intraepithelial lymphocytes requires occludin. Proc. Natl. Acad. Sci. 109, 7097–7102.

Edelblum, K. L., Singh, G., Odenwald, M. A., Lingaraju, A., El Bissati, K., McLeod, R., Sperling A.I. & Turner, J. R. (2015). γδ intraepithelial lymphocyte migration limits transepithelial pathogen invasion and systemic disease in mice. Gastroenterology, 148, 1417–1426.

Fischer, M. A., Golovchenko, N. B., & Edelblum, K. L. (2020). γδ T cell migration: Separating trafficking from surveillance behaviors at barrier surfaces. Immunol. Rev. 298, 165–180.

Foucault, I., Le Bras, S., Charvet, C., Moon, C., Altman, A., & Deckert, M. (2005). The adaptor protein 3BP2 associates with VAV guanine nucleotide exchange factors to regulate NFAT activation by the B-cell antigen receptor. Blood, 105, 1106–1113.

Girón-Pérez, D. A., Vadillo, E., Schnoor, M., & Santos-Argumedo, L. (2020). Myo1e modulates the recruitment of activated B cells to inguinal lymph nodes. J. Cell Sci., 133, jcs235275.

Girón-Pérez, D. A., Piedra-Quintero, Z. L., & Santos-Argumedo, L. (2019). Class I myosins: Highly versatile proteins with specific functions in the immune system. J. Leukoc. Biol. 105, 973–981.

Goodman, T., & Lefrançois, L. (1988). Expression of the γ-δ T-cell receptor on intestinal CD8+ intraepithelial lymphocytes. Nature, 333, 855–858.

He, S., Kahles, F., Rattik, S., Nairz, M., McAlpine, C. S., Anzai, A., Selgrade D., Fenn A.M., Chan C. T., Mindur J. E., et al. (2019). Gut intraepithelial T cells calibrate metabolism and accelerate cardiovascular disease. Nature, 566, 115–119.

Hensel, A., Stahl, P., Moews, L., König, L., Patwardhan, R., Höing, A., Schulze N., Nalbant P., Stauber R. H. & Knauer, S. K. (2022). The Taspase1/Myosin1f-axis regulates filopodia dynamics. Iscience, 25, 104355.

Hu, M. D., Ethridge, A. D., Lipstein, R., Kumar, S., Wang, Y., Jabri, B., Turner J. R. & Edelblum, K. L. (2018). Epithelial IL-15 is a critical regulator of γδ intraepithelial lymphocyte motility within the intestinal mucosa. J. Immunol. 201, 747–756.

Hu, M. D., Golovchenko, N. B., Burns, G. L., Nair, P. M., Kelly IV, T. J., Agos, J., Irani M. Z., Soh W. S., Zeglinski M. R., Lemenze A., et al. (2022). γδ intraepithelial lymphocytes facilitate pathological epithelial cell shedding via CD103-mediated granzyme release. Gastroenterology, 162, 877–889.

Kim, S. V., Mehal, W. Z., Dong, X., Heinrich, V., Pypaert, M., Mellman, I., Dembo M., Mooseker M. S., Wu D. & Flavell, R. A. (2006). Modulation of cell adhesion and motility in the immune system by Myo1f. Science, 314, 136–139.

Krendel, M., & Mooseker, M. S. (2005). Myosins: tails (and heads) of functional diversity. Physiology, 20, 239–251.

Krendel, M., Kim, S. V., Willinger, T., Wang, T., Kashgarian, M., Flavell, R. A., & Mooseker, M. S. (2009). Disruption of Myosin 1e promotes podocyte injury. J. Am. Soc. Nephrol. 20, 86–94.

López-Ortega, O., & Santos-Argumedo, L. (2017). Myosin 1g contributes to CD44 adhesion protein and lipid rafts recycling and controls CD44 capping and cell migration in B lymphocytes. Front. Immunol. 8, 1731.

López-Ortega, O., Ovalle-García, E., Ortega-Blake, I., Antillón, A., Chávez-Munguía, B., Patiño-López, G., Fragoso-Soriano R. & Santos-Argumedo, L. (2016). Myo1g is an active player in maintaining cell stiffness in B-lymphocytes. Cytoskeleton, 73, 258–268.

Maravillas-Montero, J. L., & Santos-Argumedo, L. (2012). The myosin family: unconventional roles of actin-dependent molecular motors in immune cells. J. Leukoc. Biol. 91, 35–46.

McIntyre, C. L., Monin, L., Rop, J. C., Otto, T. D., Goodyear, C. S., Hayday, A. C., & Morrison, V. L. (2020). β2 Integrins differentially regulate γδ T cell subset thymic development and peripheral maintenance. Proc. Natl. Acad. Sci. 117, 22367–22377.

Navinés-Ferrer, A., & Martín, M. (2020). Long-tailed unconventional class I myosins in health and disease. Int. J. Mol. Sci. 21, 2555

Navinés-Ferrer, A., Ainsua-Enrich, E., Serrano-Candelas, E., Proaño-Pérez, E., Muñoz-Cano, R., Gastaminza, G., Olivera A. & Martin, M. (2021). MYO1F Regulates IgE and MRGPRX2-Dependent Mast Cell Exocytosis. J. Immunol. 206, 2277–2289.

Navinés-Ferrer, A., Ainsua-Enrich, E., Serrano-Candelas, E., Sayós, J., & Martin, M. (2019). Myo1f, an unconventional long-tailed myosin, is a new partner for the adaptor 3BP2 involved in mast cell migration. Front. Immunol. 10, 1058.

Piedra-Quintero, Z. L., Serrano, C., Villegas-Sepúlveda, N., Maravillas-Montero, J. L., Romero-Ramírez, S., Shibayama, M., Medina-Contreras O., Nava P. & Santos-Argumedo, L. (2019). Myosin 1F regulates M1-polarization by stimulating intercellular adhesion in macrophages. Front. Immunol. 9, 3118.

Qiu, Z., & Sheridan, B. S. (2018). Isolating lymphocytes from the mouse small intestinal immune system. JoVE. 132, e57281

Salvermoser, M., Pick, R., Weckbach, L. T., Zehrer, A., Löhr, P., Drechsler, M., Sperandio M., Soehnlein O. & Walzog, B. (2018). Myosin 1f is specifically required for neutrophil migration in 3D environments during acute inflammation. Blood, 131, 1887–1898.

Schön, M. P., Arya, A., Murphy, E. A., Adams, C. M., Strauch, U. G., Agace, W. W., Donohue J.P., Her H., Beier D. R., Olson S., et al. (1999). Mucosal T lymphocyte numbers are selectively reduced in integrin αE (CD103)-deficient mice. J. Immunol. 162, 6641–6649.

Simons, K., & Toomre, D. (2000). Lipid rafts and signal transduction. Nat. Rev. Mol. Cell Biol. 1, 31–39.

Sit, S. T., & Manser, E. (2011). Rho GTPases and their role in organizing the actin cytoskeleton. J. Cell Sci. 124, 679–683.

Sumida, H., Lu, E., Chen, H., Yang, Q., Mackie, K., & Cyster, J. G. (2017). GPR55 regulates intraepithelial lymphocyte migration dynamics and susceptibility to intestinal damage. Sci. Immunol. 2, eaao1135.

Sun, W., Ma, X., Wang, H., Du, Y., Chen, J., Hu, H., Gao R., He R., Peng Q., Cui Z., et al. (2021). MYO1F regulates antifungal immunity by regulating acetylation of microtubules. Proc. Natl. Acad. Sci. 118.

Swamy, M., Abeler-Dörner, L., Chettle, J., Mahlakõiv, T., Goubau, D., Chakravarty, P., Ramsay G., Sousa C. R., Staeheli P., Blacklaws B. A., et al. (2015). Intestinal intraepithelial lymphocyte activation promotes innate antiviral resistance. Nat. Commun. 6, 1–12.

Tinevez, J. Y., Perry, N., Schindelin, J., Hoopes, G. M., Reynolds, G. D., Laplantine, E., Bednarek S. Y., Shorte S. L. & Eliceiri, K. W. (2017). TrackMate: An open and extensible platform for single-particle tracking. Methods, 115, 80–90.

Uehara, S., Grinberg, A., Farber, J. M., & Love, P. E. (2002). A role for CCR9 in T lymphocyte development and migration. J. Immunol. 168, 2811–2819.

Vadillo, E., Chánez-Paredes, S., Vargas-Robles, H., Guerrero-Fonseca, I. M., Castellanos-Martínez, R., García-Ponce, A., Nava P., Girón-Pérez D. A., Santos-Argumedo L. & Schnoor, M. (2019). Intermittent rolling is a defect of the extravasation cascade caused by Myosin1e-deficiency in neutrophils. Proc. Natl. Acad. Sci. 116, 26752–26758.

van Konijnenburg, D. P. H., Reis, B. S., Pedicord, V. A., Farache, J., Victora, G. D., & Mucida, D. (2017). Intestinal epithelial and intraepithelial T cell crosstalk mediates a dynamic response to infection. Cell, 171, 783–794.

Wagner, N., Löhler, J., Kunkel, E. J., Ley, K., Leung, E., Krissansen, G., Rajewsky K. & Müller, W. (1996). Critical role for β7 integrins in formation of the gut-associated lymphoid tissue. Nature, 382, 366–370.

Wang, X., Sumida, H., & Cyster, J. G. (2014). GPR18 is required for a normal CD8αα intestinal intraepithelial lymphocyte compartment. J. Exp. Med. 211, 2351–2359.

Wang, Y., Jin, H., Wang, W., Wang, F., & Zhao, H. (2019). Myosin1f-mediated neutrophil migration contributes to acute neuroinflammation and brain injury after stroke in mice. J. Neuroinflammation, 16, 1–9.

Wenzel, J., Ouderkirk, J. L., Krendel, M., & Lang, R. (2015). Class I myosin Myo1e regulates TLR 4-triggered macrophage spreading, chemokine release, and antigen presentation via MHC class II. Eur. J. Immunol. 45, 225–237.

Wurbel, M. A., Malissen, M., Guy-Grand, D., Malissen, B., & Campbell, J. J. (2007). Impaired accumulation of antigen-specific CD8 lymphocytes in chemokine CCL25-deficient intestinal epithelium and lamina propria. J. Immunol. 178, 7598–7606.

Wurbel, M. A., Malissen, M., Guy-Grand, D., Meffre, E., Nussenzweig, M. C., Richelme, M., Carrier A. & Malissen, B. (2001). Mice lacking the CCR9 CC-chemokine receptor show a mild impairment of early T-and B-cell development and a reduction in T-cell receptor γδ+ gut intraepithelial lymphocytes. Blood, 98, 2626–2632.

